# Assessment of the pharmacological and safety profile of the small molecule MP-004 after topical eye drops administration

**DOI:** 10.1101/2025.10.09.681374

**Authors:** Araceli Lara-López, María Rodríguez-Hidalgo, Miren Sarasola-Gastesi, Klaudia Gonzalez-Imaz, Julián Zayas, Maialen Sagartzazu-Aizpurua, José Ignacio Miranda, Ana Espinosa, Adolfo López de Munain, Jesús María Aizpurua, Francisco Javier Gil-Bea, Ainara Vallejo-Illarramendi, Javier Ruiz-Ederra

**Affiliations:** Miramoon Pharma, S.L., 20014 Donostia-San Sebastian, Spain; Groups of Sensorial Neurodegeneration and Neuromuscular Diseases, Neuroscience Area, Biogipuzkoa Health Research Institute (IIS Biodonostia), 20014 Donostia-San Sebastian, Spain; Group of Neurosciences, Departments of Pediatrics and Neuroscience, Faculty of Medicine and Nursing, University of Basque Country (UPV/EHU), 20014 Donostia-San Sebastian, Spain; Department of Dermatology, Ophthalmology and ORL, University of Basque Country (UPV/EHU), 20014 Donostia-San Sebastian, Spain; CIBERNED, ISCIII (CIBER, Carlos III Institute, Spanish Ministry of Sciences and Innovation), 28031, Madrid, Spain; Department of Organic Chemistry-I, Korta Research Center, University of the Basque Country (UPV/EHU), 20018 Donostia-San Sebastian, Spain; Department of Neurology, Hospital Universitario Donostia. OSAKIDETZA, 20014 Donostia-San Sebastián, Spain; Centro de Investigación Biomédica en Red Sobre Enfermedades Neurodegenerativas (CIBERNED), Madrid, Spain; IKERBASQUE Basque Foundation for Science, Bilbao, Spain; Department of Health Sciences, Public University of Navarra, Pamplona, Spain

**Keywords:** Small molecule, IRD, Eye drop, Pharmacokinetics, Pharmacodynamics, Toxicology

## Abstract

MP-004 is a novel small molecule under development as a non-invasive therapeutic for inherited retinal dystrophies (IRDs), including retinitis pigmentosa. Here, we evaluated its ocular concentration and safety profile following topical administration in three animal species —mouse, rabbit, and pig—. MP-004 formulation with 0.3% hyaluronic acid significantly enhanced retinal concentration in mice compared to non-formulated compound (6.67 vs 1.27 µg/g at 4 h). MP-004 reached therapeutically relevant retinal concentrations in all species tested, with minimal systemic exposure: in mice, levels detected in serum were lower than 5 ng/mL while in rabbits, the compound was undetectable in blood and peripheral tissues 12 h post-dose.

Seven-day repeated-dose toxicity studies in mice and rabbits showed no systemic or ocular toxicity. In rabbits, only transient ocular redness was observed post-administration, with no evidence of corneal damage or systemic adverse effects. Hematological and biochemical parameters in both species remained within normal limits. Importantly, retinal and optic nerve concentrations remained detectable in rabbits 7 days after final dosing, while the compound was eliminated systemically, supporting prolonged target tissue retention.

These findings demonstrate that MP-004 is well tolerated and achieves effective retinal concentrations via topical delivery, overcoming a major barrier in retinal drug development. The favorable pharmacokinetic and safety profile across species, including anatomically relevant models, supports MP-004’s advancement toward regulatory preclinical studies and clinical translation as a non-invasive therapy for IRDs.

## Introduction

MP-004 belongs to the MP compounds family, previously known as AHK compounds (Aizpurua et al. 2021). This group of small molecules is protected by international patents WO2017203083A1 and WO2023247712A1. These patents describe an innovative dual mechanism of action, which includes the normalization of intracellular calcium levels and the reduction of reactive oxygen species (ROS). Specifically, MP-004 compound is a highly water-soluble salt that has been optimized for ophthalmic applications and synthesized by CuAAC “click” methodology (Passannante et al. 2023). MP-004 has been demonstrated to be both efficacious *in vitro* and *in vivo* in models of retinitis pigmentosa (Lara-López et al. 2025).

Inherited retinal dystrophies (IRDs) are a group of neurodegenerative disorders that are commonly caused by mutations, leading to vision loss due to progressive degeneration of photoreceptors. These diseases are not amenable to effective treatment, and collectively they have a prevalence of approximately 1 in 3500 individuals. Among these conditions, retinitis pigmentosa (RP) is characterized by rod degeneration, and is the leading cause of inherited visual impairment, affecting 1 in 4000 individuals (Suleman 2025).

To date, more than 332 genes associated with IRDs have been identified according to the Retinal Information Network (RetNet, https://RetNet.org/). Despite the extensive number of genes associated with these diseases, the underlying cause of these pathologies remains to be elucidated with precision (Mustafi et al. 2019). The mechanisms undelaying cell death are contingent, at least in part, on the genetic mutation that triggers the disease. However, a combination of factors, including dysregulation of calcium homeostasis, imbalance of cGMP levels, reticulum stress and oxidative stress, is frequently observed (Bighinati et al. 2024; Manley et al. 2023; Newton and Megaw 2020; Power et al. 2020; Ziccardi et al. 2019). As demonstrated by Ma and collaborators, the ryanodine receptor (RyR), particularly the RyR2 isoform, plays a pivotal role in calcium dysregulation within photoreceptors (Ma et al. 2019). The functional state of the RyR channel is regulated by the prolyl-isomerase FKBP12, which functions as a key modulator of RyR channel gating. MP-004 has been shown to enhance the interaction between FKBP12/RyR, a mechanism implicated in calcium homeostasis and potentially relevant to pathological processes (Lara-López et al. 2025).

Oxidative stress and excessive production of reactive oxygen species are well-established contributors to the pathophysiology of IRDs, as demonstrated in several animal models (Haruta et al. 2009). Given the excellent efficacy results previously obtained with MP-004, this study aims to characterize its pharmacological properties to support its development as a new treatment for IRDs.

## Materials and Methods

### In vitro studies

Pharmacodynamics (PD) was analyzed by an *in vitro* screening of MP-004 against a standardized safety panel to assess potential off-target interactions and adverse effects. Pharmacokinetics (PK) profiling included assessment of permeability, plasma protein binding, metabolite identification, cytochrome P450 enzyme inhibition using liver microsomes, plasma stability, dissociation constant, aqueous solubility and partition coefficient. Cardiac safety pharmacology to determine MP-004 effect on hERG current was also evaluated. Detailed protocols for each assay are provided in the Supplementary Materials. *In vitro studies* were conducted by WuXi AppTec, Vivotecnia or ERBC.

### In vivo studies

#### MP-004 concentration determination assays

To assess ocular and systemic MP-004 concentrations-curves over time, the compound was administered via ocular instillation in *C57BL6/J* mice, *Dutch Belted* rabbits, and pigs, followed by euthanasia at predetermined time points. Mice received 3 μL of 20 mM MP-004 (equivalent to 18 μg/eye), rabbits received 50 μL of 40 mM MP-004 (558 μg/eye) and pigs received 100 μL of 20 mM MP-004 (558 μg/eye). Following euthanasia, blood was collected, eyes were enucleated, and retina, optic nerve and vitreous tissues were carefully dissected. In rabbits, additional systemic samples —brain, liver and kidney— were also collected to evaluate systemic distribution.

#### Seven-day repeated-dose toxicity studies

##### Mice

Seven groups of *C57BL6/J* mice (n=6 animals per group; 3 males and 3 females) were used to evaluate the ocular and systemic toxicity of MP-004 following repeated administration. Mice were anesthetized with 2% isoflurane via inhalation, and 3 µL of MP-004 (35 µg) was instilled into the right eye, while the left eye received 3 µL of vehicle (0.3% hyaluronic acid) as a control. Treatments were administered once daily for seven consecutive days. Mice were euthanized at 1, 24, 48 and 72 hours, and at 5 and 7 days following the final dose. Euthanasia was performed via exsanguination by cardiac puncture to obtain blood samples, followed by enucleation of the eyes and dissection of organs. Tissues collected included the eyes, blood, brain, liver, kidney, heart, spleen and thymus. Those were fixed in formalin at room temperature for 48 h, then transferred to phosphate-buffered saline (PBS) and stored at 4°C until histological processing at the Complutense University of Madrid. Blood samples were submitted to the Donostia University Hospital for biochemistry and hematological analyses.

##### Rabbit

Three groups of *New Zealand White* rabbits (n=4 animals per group; 2 males and 2 females) were used to assess ocular and systemic toxicity of MP-004 following repeated topical administration. A 35 µL drop containing 2.5 mg MP-004 or vehicle (0.3% hyaluronic acid) was instilled into both eyes once daily for seven consecutive days. Rabbits were euthanized 1 h and 7 days after the final dose for toxicological evaluation. Rabbits were anesthetized with intramuscular ketamine and xylazine, followed by euthanized via CO_2_ asphyxiation. Blood samples were collected via cardiac puncture. Eyes, blood, brain, liver, kidneys, heart, spleen and thymus were collected, fixed in formalin for 48 h and then transferred to PBS and stored at 4°C until histopathological processing at the Complutense University of Madrid. Biochemical and hematological analysis were performed at Donostia University Hospital.

Prior to euthanasia, all animals underwent fluorescein staining to assess corneal surface integrity. Fluorescein was applied topically, and eyes were examined under ultraviolet light to detect corneal epithelial defects.

#### Tissue processing for HPLC-MS/MS analysis

Tissue samples were mechanically pulverized in liquid nitrogen using a steel mortar pre-cooled on dry ice. Mouse retinas were immersed in 50 µl of 0.9% sodium chloride (NaCl), subjected to multiple freeze-thaw cycles, and homogenized using a mechanical tissue disruptor followed by a 30-second ultrasonic bath. A precipitating agent (1% formic acid in acetonitrile) was then added to the pulverized tissue in a volume equivalent to the sample mass for tissue samples, and three times the sample volume for serum and mouse retina samples. The mixtures were subsequently sonicated and centrifuged at 9,600xg for 5 minutes at 4°C. Supernatants were collected and analyzed by high-performance liquid chromatography-tandem mass spectrometry (HPLC-MS/MS) at the SGIker Research General Services facility, UPV/EHU (Vitoria-Gasteiz, Spain).

## Results

### In vitro studies

The secondary pharmacodynamics of MP-004 were evaluated using an *in vitro* safety panel comprising 44 selected protein targets. MP-004 exhibited no significant off-target activity, with inhibition values below 35% at concentration of 10 µM (Supplementary Figure 1).

The *in vitro* pharmacokinetic properties of MP-004 were assessed, focusing on Absorption, Distribution, Metabolism and Excretion (ADME) parameters. For absorption, MP-004 demonstrated high permeability in Caco-2 cell monolayers, with an apparent permeability coefficient (P_app_) of 40.2 ×10^-6^ cm/s in the apical-to-basolateral (A→B) direction and 31.8 ×10^-6^ cm/s in the basolateral-to-apical (B→A) direction. These values exceeded the established high permeability threshold (P_app_ ≥ 2.5 ×10^-6^ cm/s) (Supplementary Figure 2). Moreover, MP-004 showed an efflux ratio below 2 without transporter inhibitors, suggesting it is not significantly transported by P-glycoprotein (P-gp) in Caco-2 cells (Supplementary Figure 3).

In distribution studies, MP-004 showed moderate plasma protein binding across multiple species, including CD-1 mouse, Sprague-Dawley rat, New Zealand White rabbit, Beagle dog, Göttingen minipig, Cynomolgus monkey, and human. Plasma protein binding values ranged from 49.8% to 66.7%, with 62.25% binding observed in human plasma (Supplementary Figure 4).

Metabolic profiling in liver microsomes from mouse, rat, dog, rabbit, minipig and human, revealed ten distinct MP-004 metabolites. The primary metabolic pathways were mono-oxidation and demethylation (Supplementary Figure 5). MP-004 showed low potential for cytochrome P450 (CYP) inhibition, with IC_50_ values exceeding 35 µM for all isoforms tested (1A2, 2C9, 2C19, 2D6 and 3A4) (Supplementary Figure 6). In addition, MP-004 was stable in human plasma, with 99.5% of the compound remaining after incubation and a half-life (T_1/2_) exceeding 289.1 minutes (Supplementary Figure 7).

Some physicochemical parameters of MP-004 were studied. MP-004 had a mean dissociation constant (pKa) of 9.17 (Supplementary Figure 8). It was classified as a soluble compound, with a mean kinetic solubility of 74 μg/mL (Supplementary Figure 9). The compound exhibited a favorable balance between permeability and solubility, with a partition coefficient (log P) of 1.77. The distribution coefficient (log D) at physiological pH (7.4) was 1.49, indicating moderate solubility and permeability (Supplementary Figure 10).

Finally, safety pharmacology assessment using the patch clamp technique was conducted to evaluate the potential cardiac toxicity of MP-004. MP-004 induced a dose-dependent inhibition of the hERG tail current at concentrations of 9×10⁻⁷ M, 9×10⁻⁶ M, and 9×10⁻⁵ M, resulting in 4% ± 1%, 10% ± 1%, and 53% ± 5% inhibition, respectively (Supplementary Figure 11). Given the concentration-dependent effect and the fact that the mean maximum inhibition exceeded 30% at the highest concentration, an IC₅₀ value for hERG tail current inhibition was calculated by curve fitting for each individual cell. The mean IC₅₀ value derived from three individual measurements was 84 ± 18 µM. Under identical experimental conditions, the reference compound E-4031 (tested at 1, 10, and 100 nM) produced a robust inhibition of the hERG tail current (91% at the highest concentration), with an IC₅₀ of 23 nM, thereby supporting the conclusion that MP-004 exhibits minimal hERG inhibition.

### MP-004 concentration determination assays

The ocular and systemic concentration of MP-004 was assessed in mice, rabbits, and pigs following topical ocular administration.

#### In Mice

MP-004 concentration in the retina and serum was evaluated at multiple time points (2, 4, 8, 12, 24, 36 and 48 h) following a single ocular dose of MP-004 (18 µg/eye) using two vehicles (Figure 1). In the retina, MP-004 was detected up to 48 h post-administration with both vehicles analyzed. The non-formulated vehicle (PBS) showed MP-004 concentrations of 1.19 µg/g, 48 h post-administration. When formulated in 0.3% hyaluronic acid, 1.01 µg/g of MP-004 was detected in the retina at 48 h. The formulated MP-004 concentrations (Figure 1B) were higher than the non-formulated ones (Figure 1A) during most of the timepoints analyzed in the study. In fact, the MP-004 retinal concentration at 24 h post-administration with PBS (1.02 µg/g), was equivalent to the concentration observed at 48 h with the hyaluronic acid formulation, suggesting that the formulation enhances the residence time of MP-004 in the target tissue.

**Figure 1.**
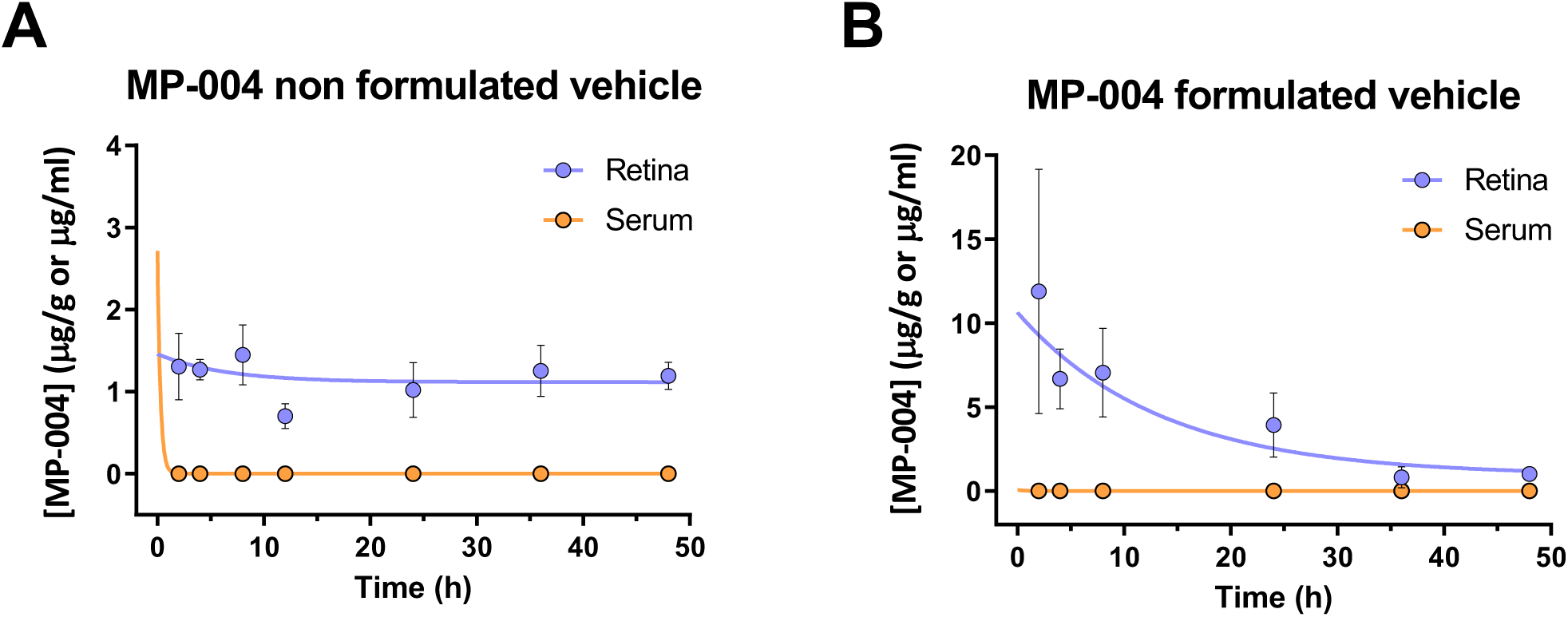
Ocular and systemic MP-004 concentrations-curves in mice after topical administration with a non-formulated (A) and formulated (B) vehicle. The retina availability of the MP-004 compound was studied in retina for 48 h. Both non-formulated MP-004 (vehicle PBS) and formulated MP-004 (vehicle 0.3% hyaluronic acid) were detected in retina up to 48 h after administration, being the formulated one higher than non-formulated. In serum the non-formulated MP-004 was detected 2 h after-dose but not at later times. MP-004 formulated with the 0.3% hyaluronic acid vehicle was detected in serum up to 4 h after administration. Data are expressed as mean ± SEM. n=3-8 mice/time.

In serum, MP-004 was detected at 2 h post-administration when delivered with the PBS vehicle (Figure 1A), reaching a concentration of 1.98 ng/ml. It subsequently fell below the limit of quantification and reappeared at 36 and 48 h, with concentrations of 0.14 and 0.13 ng/ml, respectively. In contrast, when administered with the hyaluronic acid vehicle (Figure 1B), MP-004 remained detectable in serum for a longer duration. At 4 h post-administration, a concentration of 0.27 ng/ml was detected. After that point, levels fell below the limit of quantification and remained undetectable through 48 h.

The area under the concentration–time curve (AUC) of the formulated vehicle was 185.18 μg/g·h for the retina and 0.009 μg·ml/·h for the serum. Based on these values, the retina-to-serum AUC ratio was calculated to be approximately 0.005%, indicating minimal systemic exposure and suggesting that ocular instillation results in a very low drug availability in the serum.

#### In Rabbits

MP-004 concentration was assessed at 0.5, 1, 2, 4, 8, 12, 24, 48 and 72 h post-administration of 585 µg/eye of MP-004 formulated in 0.3% hyaluronic acid. Samples were collected from ocular tissues —retina and optic nerve—, extraocular tissues — brain, liver and kidney—, and blood —serum—. MP-004 was detected in all tissues analyzed (Figure 2).

– Ocular tissues (Figure 2A): retinal concentrations were the highest, with detectable levels in the retina up to 48 h post-administration. In the optic nerve, peak concentration was observed at 1 h, followed by a progressive decline and complete clearance by 24 h.
– Extraocular tissues (Figure 2B): Lower MP-004 levels were detected in brain, kidney, and liver, with no detectable compound beyond 8 h post-dose.
– Serum (Figure 2C): MP-004 presented the maximum concentration at 0.5 h (43.6 ng/ml), subsequently decreasing until be undetectable 4 h post-dose.

**Figure 2.**
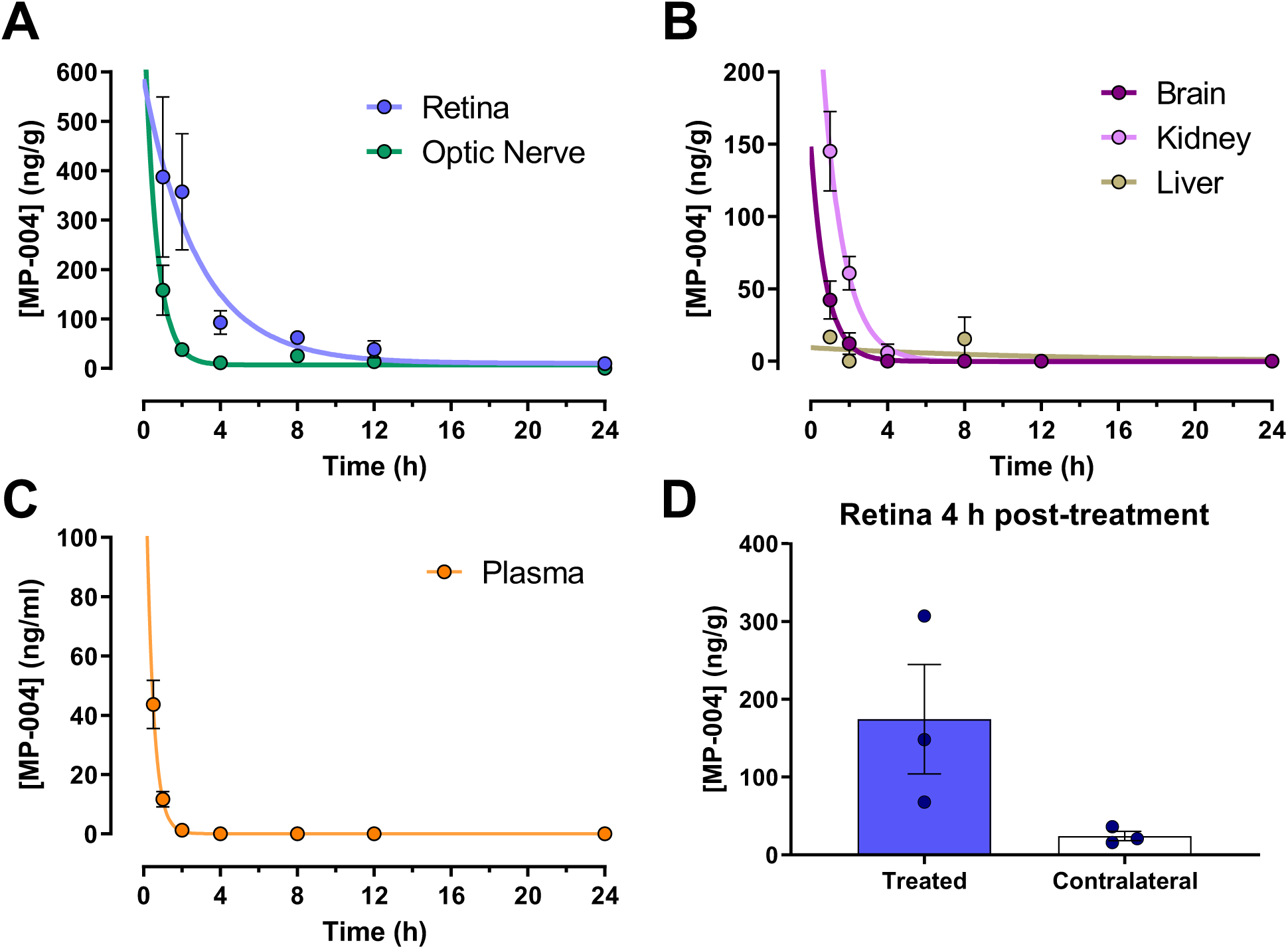
MP-004 pharmacokinetics after topical ocular administration in rabbit. MP-004 concentration in (**A**) ocular —retina and optic nerve— and (**B**) extraocular — brain, kidney, liver— and (**C**) plasma tissues. **D)** Concentration of MP-004 in the treated and contralateral (untreated) eye. Data are expressed as mean ± SEM. n=3 rabbits/time.

Importantly, bilateral distribution was observed: following unilateral ocular administration, MP-004 was detected in both retinas at 4 h post-treatment. The concentration in the contralateral (untreated) eye was approximately seven-fold lower than in the treated eye (174.3 vs. 15.8 ng/g, Figure 2D).

Pharmacokinetic parameters were calculated for rabbits (Supplementary Figure 12):

– In serum, the maximum concentration (C_max_) was 43.6 ng/ml at 0.5 h and 12.3 ng/ml at 1 h, with a half-life (T_1/2_) of around 20 minutes and T_max_ at 0.5 h. The area under the serum concentration-time curve from time zero to the last quantifiable concentration (AUC_0-last_) value was 27.7 ng·h/ml.
– In retina, the C_max_ was 387 ng/g at 1 h, with a T_1/2_ of 6.34 h and AUC_₀–last_ of 1460 ng·h/g. This corresponds to a retina-to-serum concentration ratio of 32 at 1 h and 158 at 2 h.
– For extraocular tissues, C_max_ values were 58 ng/g (optic nerve), 145 ng/g (kidney), 42.2 ng/g (brain), and 16.7 ng/g (liver), all peaking at 1 h and undetectable by 8 h post-dose.

#### In Pigs

MP-004 tissue distribution was evaluated 4 h after ocular administration of 585 µg/eye in 0.3% hyaluronic acid. The compound was detected in all tissues examined (retina, optic nerve, vitreous and serum), with the highest concentration in the retina (20.61 ng/g), and the optic nerve (31.12 ng/g). A low concentration (2.84 ng/g) was detected in serum (Figure 3).

**Figure 3.**
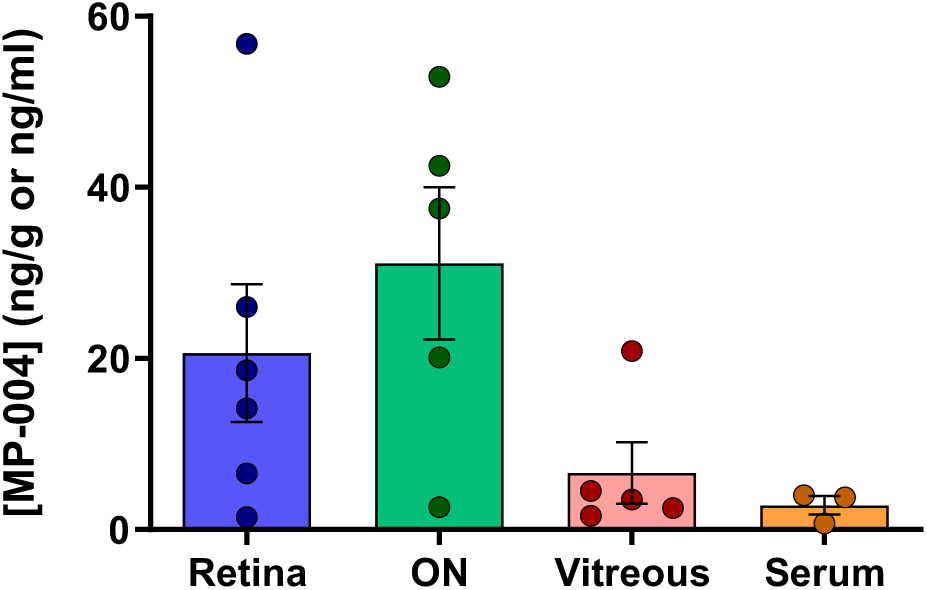
MP-004 availability in pig after ocular topical administration. MP-004 concentration in ocular tissues —retina, optic nerve (ON), vitreous— are expressed as ng of MP-004/g of tissue. MP-004 serum concentration is expressed as ng of MP-004/ml of serum. Data are expressed as mean ± SEM. n=6 eyes.

### Seven days repeated-dose toxicity studies

#### In Mice

The maximum feasible dose (MFD) of MP-004 in mice was 35 µg/eye/day. Animals received daily topical ocular instillation in one eye for 7 consecutive days. MP-004 was well tolerated throughout the study period, and no clinical adverse effects were observed. Animals were sacrificed at various time points post-final administration (1, 24, 48 and 72 h, and 5 and 7 days) for histopathological and blood analyses.

Histopathological ocular examination revealed no remarkable pathological findings in any of the treated animals (Figure 4). Minor findings —such as a mild lymphoid infiltrate in the renal pelvis or multifocal subacute pneumonia in the lungs— were observed sporadically across both treated and control groups. No differences were detected between time points, between treated and untreated groups, or between treated and contralateral (vehicle-treated) eyes Biochemical parameters (Supplementary Figure 13) were largely within normal ranges. However, a transient and statistically significant decrease in serum chloride concentrations was observed at 24, 48, and 72 h post-administration (p=0.0394, p=0.0163 and p=0.0104, respectively), with a nadir of 109.7 mEq/L at 72 h, compared to 115 mEq/L in untreated controls. By day 5, chloride levels returned to baseline (112.7 mEq/L on day 7). Liver markers aspartate aminotransferase (AST) and alanine aminotransferase (ALT) showed a slight, non-significant increase on day 7, but values remained within the range observed in controls, supporting the absence of hepatic toxicity.

**Figure 4.**
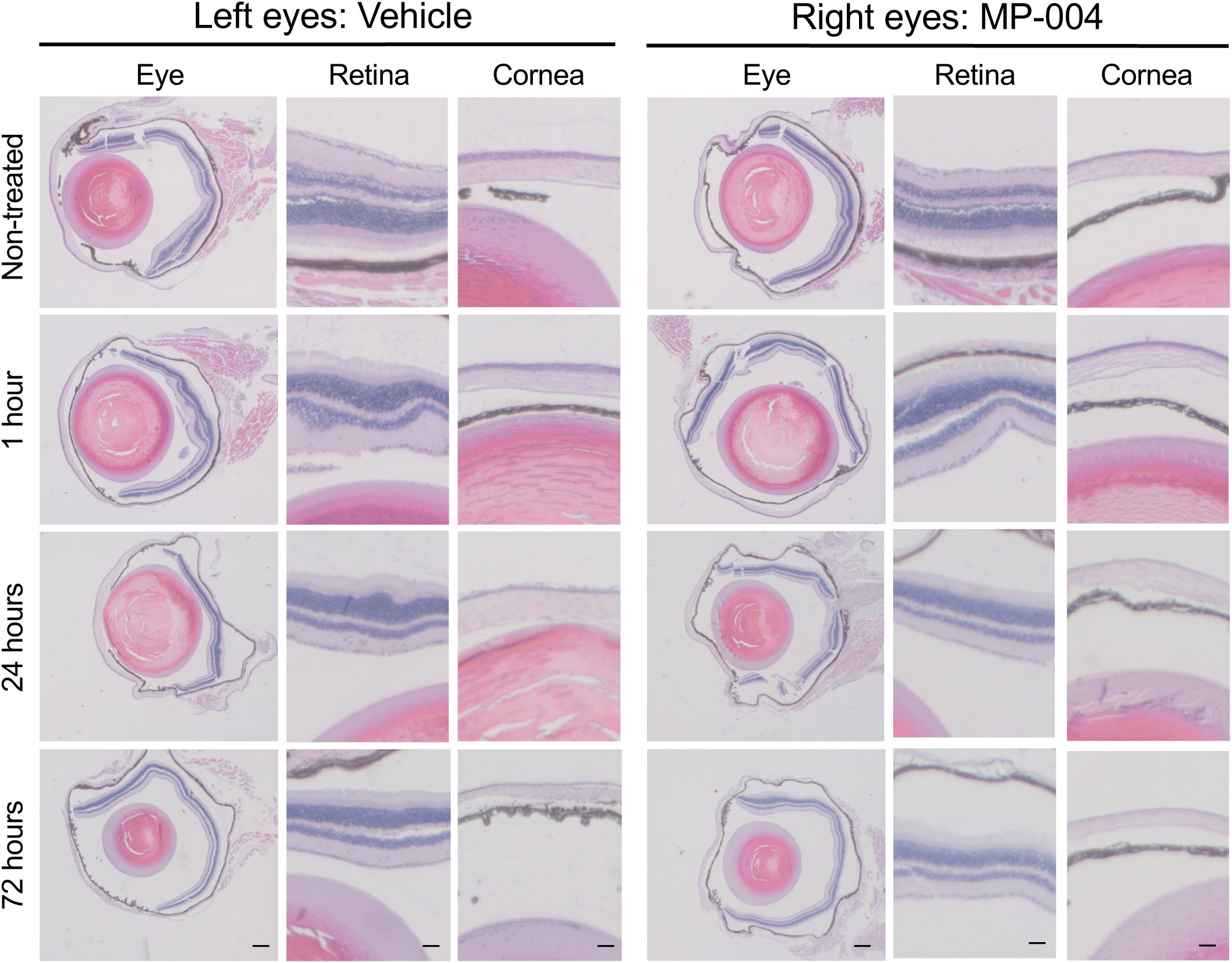
Representative images from histopathological analysis of mice eyes. Left panel shows images of left eyes (vehicle-treated) and right panel of right eyes (MP-004-treated). Images of the non-treated, 1, 24 and 72 h post-treatment groups are shown. Scale bar: Eyes images: 200 μm. Retina and cornea images: 50 μm.

Hematological analysis (Supplementary Figure 14) showed no significant alterations in red or white blood cell parameters, including erythrocyte count, hemoglobin, hematocrit, reticulocytes, leukocytes, or platelets. A transient reduction in mean corpuscular volume (MCV) was observed at 24 h (52.93 fL vs. 57.87 fL in controls), persisting through day 7. Mean corpuscular hemoglobin concentration (MCHC) showed a slight increase by day 7 (26.37 vs. 24.73 g/dL). These changes were within physiological ranges and not accompanied by other hematologic abnormalities, supporting the absence of systemic hematological toxicity.

#### In Rabbits

Rabbits were administered MP-004 daily at the MFD (2.5 mg/eye/day) via ocular instillation for 7 days. A transient conjunctival redness was observed post-administration, lasting 2-5 minutes, and fully resolved within 10 minutes. Eye examinations conducted at 5, 10, and 60 minutes post-dose confirmed full recovery (Supplementary Figure 15A). Fluorescein staining performed prior to euthanasia revealed no corneal damage in either treated or control animals (Supplementary Figure 15B).

Histopathological evaluation revealed no significant ocular or systemic lesions attributable to MP-004 (Figure 5). No differences were observed between treated and control groups. Similarly, biochemical (Supplementary Figure 16) and hematological (Supplementary Figure 17) profiles showed no significant deviations. All measured parameters were within the reference range and consistent with published data (Hewitt et al. 1989; McGill 2016; Shousha, Mahmoud, and Hameed 2017).

**Figure 5.**
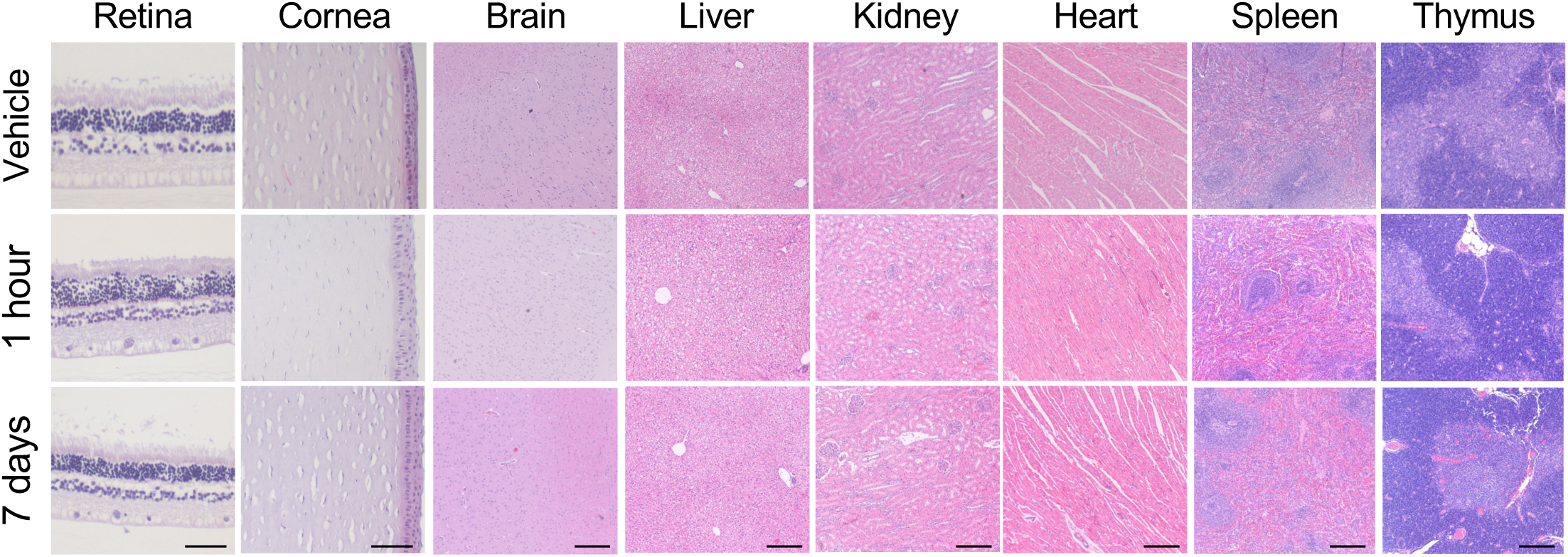
Histopathological analysis of rabbit tissues. Representative images of ocular —retina and cornea— and extraocular —kidney, heart, liver, spleen, thymus and brain— tissues of 1 h and 7 days post-dose groups are shown. Scale bar: Ocular tissues: 50 μm. Extraocular tissues: 200 μm.

MP-004 concentration analysis was assessed in rabbits following the 7-days ocular administration. Tissue distribution was analyzed in ocular —retina and optic nerve— and systemic —brain, liver, kidney and serum— compartments. In animals sacrificed 1 h after the final dose, MP-004 was detected in all analyzed tissues, with notably higher concentrations in ocular tissues compared to extraocular ones. By 7 days post-administration, MP-004 was no longer detectable in systemic tissues (Figure 6). However, MP-004 remained in the retina and optic nerve, albeit at lower concentrations: retinal levels were approximately six-fold lower (321.2 vs 2024.14 ng/g), and optic nerve levels were approximately 2.5-fold lower (734.6 vs 1801.7 ng/g) compared to those measured 1 h after the last dose. Toxicokinetic analysis in serum confirmed the presence of MP-004 1 h post-dose with no detectable levels at later time points. These results indicate that repeated ocular administration of MP-004 at the MDF results in no observable local or systemic toxicity, with systemic clearance of the compound occurring within 7 days, while sustained drug levels are maintained in the target ocular tissues — particularly the retina— over the time.

**Figure 6.**
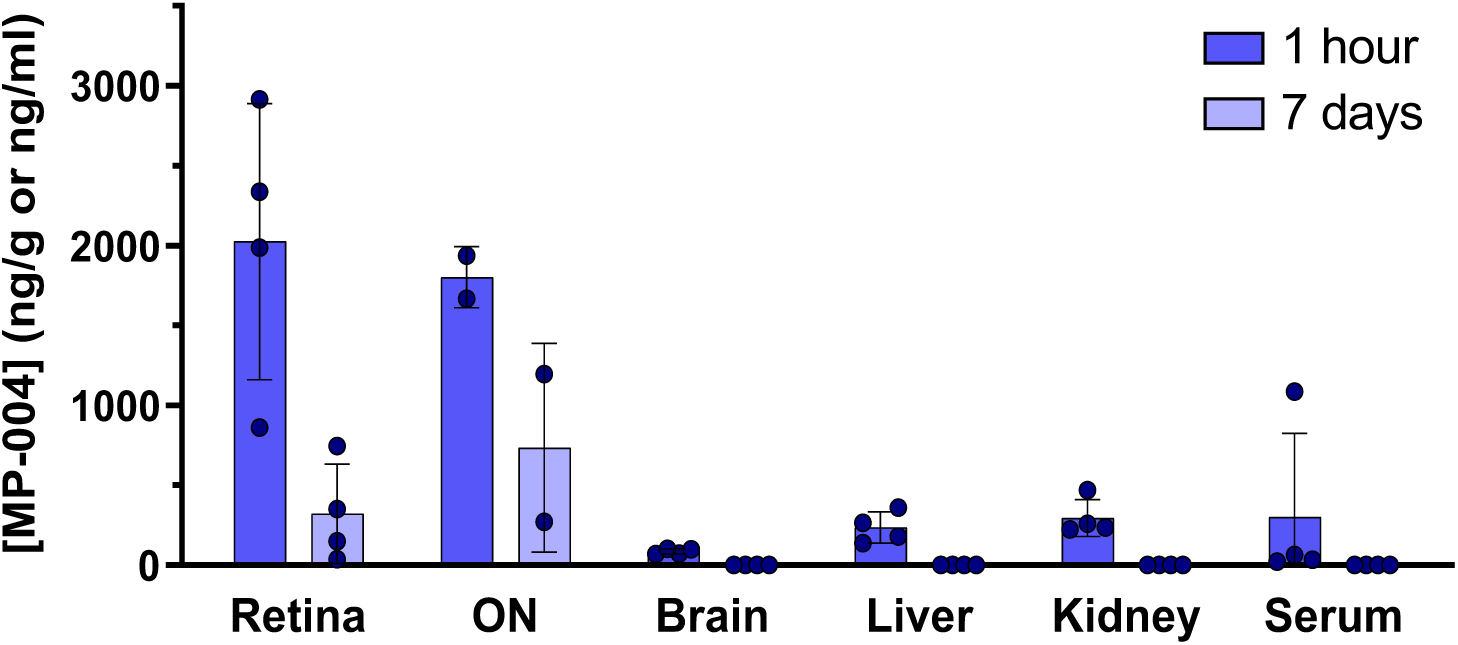
Concentration of MP-004 in the different tissues measured by HPLC-MS/MS. Availability of MP-004 in the different tissues analyzed —retina, optic nerve (ON), brain, liver, kidney— is expressed as ng of MP-004 per g of tissue. Serum concentration is expressed as MP-004/ml of serum. MP-004 was detected in all tissues analyzed 1 h after the last administration but was only detected in ocular tissues 7 days after the 7 days-treatment. Data are expressed as mean ± SEM. n=4 rabbits/time.

## Discussion

The regulatory development of a new drug candidate requires comprehensive pharmacokinetic (PK) and pharmacodynamic (PD) studies to ensure effective delivery to the target organ and to demonstrate an acceptable safety profile. In this study, we show that MP-004 exhibits a favorable safety margin, with no significant off-target activity. The cardiac safety pharmacology is also reinforced by the minimal inhibition of hERG tail current observed. Additionally, the MP-004 ADME profile supports its suitability for ocular administration in the form of eye drops. However, the effective delivery of therapeutic agents to the retina remains a major hurdle in the treatment of IRDs, particularly when non-invasive routes such as topical instillation are preferred. In this context, we evaluated the pharmacokinetics, tissue distribution, and safety profile of MP-004, a small molecule candidate with therapeutic potential for retinitis pigmentosa, across three animal models: mouse, rabbit, and pig.

Our results show that MP-004 reaches the retina efficiently after topical administration, with concentrations surpassing the effective threshold previously defined for this tissue (Lara-López et al. 2025). Importantly, the formulation vehicle played a key role in modulating tissue availability. In mice, a 0.3% hyaluronic acid formulation increased retinal concentrations at 4 h post-administration more than five-fold compared to the PBS vehicle. Moreover, the retinal concentration observed at 48 h post-dose with the hyaluronic acid formulation was comparable to that seen at 24 h with the PBS formulation, indicating improved retention in the target tissue.

Serum levels of MP-004 were low across all time points and species studied, highlighting the compound’s limited systemic exposure. In mice, peak serum concentrations at 2 h post-administration were approximately 1.98 ng/mL, several orders of magnitude lower than corresponding retinal levels. Similarly, in rabbits, MP-004 was rapidly cleared from systemic circulation and extraocular tissues, with no detectable levels in blood or peripheral organs beyond 12 h post-administration. These pharmacokinetic characteristics are advantageous for minimizing systemic toxicity and off-target effects. Furthermore, this correlates with the *in vitro* MP-004 effect on hERG tail current amplitude where the half-maximal inhibitory concentration (IC_50_) was 84 µM, which is below the maximum MP-004 concentration detected in rabbit serum (43.6 ng/ml, equivalent to 150 µM), thus confirming the safety of MP-004.

The ability of MP-004 to reach the retina following topical administration was confirmed in all three animal species. Notably, even in large-eyed animals such as rabbits and pigs—models that more closely approximate human ocular anatomy (Loiseau et al. 2023)— MP-004 was detected in the retina and optic nerve within hours of administration. These findings underscore the potential of MP-004 to overcome the traditionally limited ocular availability associated with topical therapies aimed at the posterior segment (del Amo et al. 2017; Löscher et al. 2022; Rodrigues et al. 2018).

Toxicological evaluation following 7-day repeated-dose exposure revealed no evidence of ocular or systemic toxicity in either mice or rabbits. In mice, minor changes in electrolyte levels (chloride) and erythrocyte indices (MCV and MCHC) were observed at isolated time points. These variations, however, remained within the published physiological ranges (Aldana et al. n.d.; Mazzaccara et al. 2008; Silva-Santana et al. 2020). No alterations were detected in renal or hepatic markers. Additionally, these findings were supported by the absence of histopathological findings. These changes are thus unlikely to be compound-related and may reflect transient physiological adaptations or interindividual variability (Hewitt et al. 1989; O’Connell et al. 2015; Shousha, Mahmoud, and Hameed 2017).

Similarly, in rabbits, daily administration of MP-004 at the maximum feasible dose produced only transient ocular redness that resolved within minutes post-instillation, with no signs of corneal damage or other ocular abnormalities. Hematological and biochemical parameters remained within normal ranges, and no histopathological abnormalities were observed in any major organs. Importantly, MP-004 was detectable in the retina up to 7 days after the final administration, whereas it was eliminated from systemic circulation and peripheral tissues, reinforcing the favorable target-tissue retention and clearance profile of the compound.

## Conclusions

Taken together, these findings support the continued development of MP-004 as a topically administered therapeutic for IRDs. The compound demonstrates robust retinal availability, low systemic exposure, and a favorable safety profile across species, including in anatomically relevant large animal models. These attributes, combined with the simplicity and non-invasiveness of the topical route, position MP-004 as a strong candidate for further regulatory toxicology studies and advancement toward first-in-human trials.

## List of abbreviations

ADME: Absorption, Distribution, Metabolism and Excretion
ALP: Alkaline phosphatase
ALT: Alanine aminotransferase
AST: Aspartate aminotransferase
AUC: Area under concentration-time curve from time zero to the last quantifiable concentration
CK: Creatinine kinase
Cl^-^: Chloride
C_max_: Maximum concentration
Ctrl: Control
CYP: Cytochrome P450
d: Day
h: Hour
HPLC-MS/MS: High-performance liquid chromatography-tandem mass spectrometry
IC_50_: Half-maximal inhibitory concentration
IRD: Inherited retinal dystrophies
IRF: Immature reticulocytes fraction
K: Potassium
log D: Distribution coefficient
log P: Partition coefficient
Lym: Lymphocytes
MCH: Erythrocyte mean corpuscular hemoglobin
MCHC: Mean corpuscular hemoglobin concentration
MCV: Mean corpuscular volume
MFD: Maximum feasible dose
Min: Minute
Mon: Monocytes
Na^+^: Sodium
NaCl: Sodium chloride
Neu: Neutrophils
ON: Optic nerve
PD: Pharmacodynamics
P-gp: P-glycoprotein
PK: Pharmacokinetics
pKa: Dissociation constant
RDW: Red cell distribution width
Ret: Reticulocytes
ROS: Reactive oxygen species
RP: Retinitis pigmentosa
RyR: Ryanodine receptor
T_1/2_: Half-life

## Declarations

### Availability of data and materials

The experimental datasets generated and/or analyzed during the current study are not publicly available due to presence in a proprietary data format but are available from the corresponding author upon reasonable request.

### Competing interests

A Lara-López and Ana Espinosa are employees at Miramoon Pharma S.L.; M Rodríguez-Hidalgo, None; M Sarasola-Gastesi, None; K Gonzalez-Imaz, None; Julián Zayas, None; Ana Espinosa, Employment of Miramoon Pharma S.L.; M Sagartzazu-Aizpurua, None; Francisco Javier Gil-Bea, Miramoon Pharma S.L. Founder; JI Miranda, Miramoon Pharma S.L. Founder; JM Aizpurua, Miramoon Pharma S.L. Founder; A López de Munain, Miramoon Pharma S.L. Founder; A Vallejo-Illarramendi, Miramoon Pharma S.L. Founder; Javier Ruiz-Ederra, Miramoon Pharma S.L. Founder.

### Funding

The research leading to these results was funded by MCIU/AEI/ISCIII/10.13039/501100011033 and European Union NextGenerationEU/PRTR (CPP2022-009867, OC-2024-1-27160 and PID2024-157668OB-I00 to JRE, to AVI, DIN2020-011302 to ALL); Gobierno Vasco (MTVD24/BG/003 and MTVD23/BD/008, MTVD25/BG/002 to JRE, IT1732-22 to AVI, IT1741-22 to JMA and MS-A and Pre-2019-1-0325 to MRH); Diputación Foral de Gipuzkoa (2023-CIEN-000032-01 to JRE); and grants from BEGISARE Foundation and Susana Monsma Foundation to JRE.

### Authors’ contributions

ALL, MRH, MSG, KGI and JZ substantially contributed to the experimental part of the work. MSA and JMA synthesized the MP-004 compound. ALL, AE, FGB, JIM, JMA, ALM, AVI and JRE contributed to the conception and design of the work. ALL, AVI, JRE and FGB contributed to data analysis and data interpretation. All authors subsequently contributed to drafting or revising the content.

## Acknowledgements

The quantitative determination of MP-004 in the biological samples was performed at the SGIker General Research Services of the UPV/EHU (Vitoria-Gasteiz). The author is grateful for the technical and human support provided by the SGIker of the UPV/EHU, and the European funding (ERDF and ESF). Especially to Mª Carmen Sampedro.

The authors also thank Doctors Eva Gil y Mercedes Gradin from the Hematology and Hemotherapy Laboratory of the Donostia University Hospital; and Doctor Juana Flores from the Complutense University of Madrid.

Animals’ experiments were conducted in the Animal Facility and Experimental Operating Room Platform of Biogipuzkoa HRI. Authors thanks for the technical and human support provided by the platform, especially to Elizabeth Hijona.

## Notes

### Competing Interest Statement

A.L.L. is an employee and JIM, JMA, ALdM, AVI and JRE have equity ownership in Miramoon Pharma S.L., which is developing novel triazole molecules related to the reported research. The terms of this agreement have been reviewed and approved by the University of the Basque Country and BIOEF.

