## Supplementary material for "Assessment of the pharmacological and safety profile of the small molecule MP-004 after topical eye drops administration": Supp_Material

### SUPPLEMENTAL MATERIAL

#### Materials and Methods

*In Vitro Studies*

**Safety Panel**

<u>Enzyme panel</u>:

1. **LCK (lymphocyte-specific protein tyrosine kinase)**: Compounds were transferred (100 nL) into 384-well plates using Echo acoustic dispenser. A 5 μL mixture of 40 nM enzyme and 4 μM 0%-phosphorylated Tyr2 was added to assay wells. After 30 s of 1000 rpm centrifugation, 5 μL of 180 μM ATP mix was added. Plates were incubated at 23°C for 60 min, followed by addition of 5 μL development mix (1:128 dilution in buffer), and incubated for another 60 min at 23°C. Fluorescence was read at 445 nm and 520 nm using Envision® plate reader, and the 445/520 ratio was calculated.
2. **MAO-A (monoamine oxidase A)**: Compounds (100 nL) were transferred to assay plate using Echo. 5 μL of 2X MAO substrate mixture (16 μM) was added and centrifuged (30 s, 1000 rpm). Then, 5 μL of 2X MAO-A enzyme (0.25 μM) was added, and plates were incubated at 23°C for 90 min. Detection reagent (10 μL) was subsequently added, and plates were centrifuged again and incubated for an additional 60 min at 23°C. Fluorescence was measured using a PerkinElmer Envision plate reader.
3. **PDE3A/PDE4D2 (**Phosphodiesterases**)**: Test compounds were transferred to plate. 5 μL of diluted enzyme (400 pM PDE3A and 400 pM PDE4D2) was added to each well and incubated at room temperature (RT) for 20 min. Subsequently, 5 μL of 1000 nM cAMP was added and incubated for 60 min. AMP-Glo™ Reagent I (5 μL) was added, followed by another 60-min incubation. Finally, 10 μL of AMP

detection solution was added and incubated for an additional 60 min before luminescence detection on an Envision plate reader.

4. **ACHE** (Acetylcholinesterase): TRIS-HCl buffer (50 mM, pH 8.0) and $NaH_2PO_4$–$Na_2HPO_4$ buffer (50 mM) were used to prepare assay solutions. Enzyme (4.5 nM in TRIS buffer) was dispensed (50 µL) into black 96-well plates, followed by 2 µL of test or reference compounds. Plates were shaken and incubated at RT for 5 min. A substrate mixture of 100 mM acetylthiocholine iodide and 10 mM DTNB (final concentrations 400 µM and 0.5 mM, respectively) in assay buffer was added (48 µL). Plates were centrifuged, incubated at 37°C for 1 h, and read on a SpectraMax reader.
5. **COX-1/COX-2** (Cyclooxygenases): Compounds were serially diluted in 10% DMSO, with a final DMSO concentration of 0.5%, and transferred to plates at a final concentration of 10 µM. To the plates, 100 µL of 2X COX-1 (40 nM) or 2X COX-2 (80 nM) with 250X Heme was added, and incubated at RT for 10 min. Subsequently, 50 µL each of 4X ADHP (100 µM) and 4X arachidonic acid (12 µM) were added. After centrifugation, fluorescence was measured at excitation 530 nm and emission 590 nm using a FlexStation reader.

#### Receptor and Functional Assays

FLIPR Calcium Flux Assay: Cells were seeded at 20,000 cells/well in 384-well plates. After overnight incubation at 37°C, 5% $CO_2$, cells were loaded with 20 µL of 2X Fluo-4 Direct™ No-Wash dye solution (prepared freshly with probenecid), incubated at 37°C for 50 min and RT for 10 min. Test and reference compounds were serially diluted in DMSO and transferred to compound and DRC plates. After the addition of assay buffer and centrifugation, plates were placed in the FLIPR® Tetra system. Fluorescence signals were recorded before and after compound/agonist addition. For antagonist mode, agonists were added following compound pre-incubation. $EC_{50}$ and Max–Min responses were calculated from kinetic traces.

Filtration Binding Assay: Membrane preparations and ligands were diluted to specified concentrations. Test compounds (10 μM) and control samples for total and nonspecific binding were transferred to assay. After incubation, mixtures were filtered through GF/C plates pre-soaked with 0.3% PEI. Plates were washed 4 times with cold buffer, dried at 50°C for 1 h, and sealed. Microscint 20 cocktail (50 μL) was added, and plates were sealed and read using a PerkinElmer MicroBeta2. Data were analyzed using variable slope dose–response fitting.

Nuclear Receptor Binding Assay: Cytosol and radioligand preparations were diluted to working concentrations. Test compounds (10 μM) and controls were transferred to assay plates. Cytosol (100 μL) and radioligand (100 μL) were added to each well and incubated under defined conditions. Reactions were terminated by adding 100 μL of radio-ligand adsorption buffer and shaking at 4°C for 15 min. Plates were centrifuged (4600 rpm, 30 min, 4°C), and 100 μL of supernatants were transferred to scintillation vials with 2 mL of Ultima Gold cocktail. Samples were counted using TriCap scintillation counters, and inhibition curves were fitted using nonlinear regression.

**Permeability Assay (Caco-2 cells):**

Caco-2 cells were seeded onto polyethylene membranes in 96-well Corning Insert plates at a density of 1 × $10^5$ cells/$cm^2$ and maintained with refreshed medium every 4–5 days for 21–28 days to allow formation of confluent monolayers. Bidirectional transport experiments were conducted using HBSS containing 10 mM HEPES (pH 7.4). Test compound, MP-004, and digoxin, a P-gp substrate control, were evaluated at 2 μM and 10 μM, respectively, in the presence or absence of 30 μM verapamil (a P-glycoprotein inhibitor), in bidirectional transport studies performed in duplicate. Nadolol and metoprolol were tested at 2 μM without verapamil in the A–B direction. After a 2-hour incubation at 37°C with 5% $CO_2$ and saturated humidity, samples were mixed with acetonitrile containing an internal standard and centrifuged at 3200 ×g for 10 minutes. 100 μL of the resulting supernatant were diluted with 100 μL ultrapure water and analyzed by LC-MS/MS. Compound concentrations in donor, receiver, and starting solutions were quantified based on analyte-to-internal standard peak area ratios. Monolayer integrity was confirmed post-assay by measuring lucifer yellow permeability.

**Plasma protein binding (PPB):**

Phosphate buffer (pH 7.4) was prepared by titrating a basic solution (14.2 g/L $Na_2HPO_4$ and 8.77 g/L NaCl in deionized water) with an acidic solution (15.6 g/L $NaH_2PO_4{\cdot}2H_2O$ and 8.77 g/L NaCl). The stop solution consisted of tolbutamide (200 ng/mL), labetalol (200 ng/mL), and metformin (50 ng/mL) in 100% acetonitrile. Dialysis membranes (12–14 kDa cutoff, HTDialysis) were pre-soaked in ultrapure water for 1 hour, then in 20:80 ethanol/water (v/v) for 20 minutes, and rinsed in ultrapure water before use. Plasma was thawed under cold water, centrifuged at 3220 rpm for 5 min, and pH-adjusted (7.0–8.0). Test compounds were dissolved in DMSO (10 mM stocks, diluted to 400 μM), then spiked into plasma at a 1:200 ratio. For T0 samples, 50 μL of matrix was combined with 50 μL of buffer, followed by 500 μL of stop solution, and stored at 2–8°C. In the dialysis plate, 150 μL of matrix and 150 μL of buffer were added to the donor and receiver sides, respectively, and incubated for 4 h at 37°C with 5% $CO_2$ and 100 rpm shaking. After incubation, 50 μL aliquots from both sides were collected, mixed 1:1 with the opposite matrix, and treated with 500 μL of stop solution. Samples were vortexed, centrifuged (4000 rpm, 20 min), and 100 μL of supernatant was analyzed by LC-MS/MS. Blank samples were prepared analogously using matrix-matched plasma and buffer.

**Metabolite Profiling In Liver Microsomes:**

Microsomal Incubation and Sample Processing: Microsomal stability assays were performed using liver microsomes from mouse, rat, dog, rabbit, minipig, and human at a final protein concentration of 1.12 mg/mL. The test compound and the positive control 7-ethoxycoumarin were each tested at 10 mM. Microsomal incubations (400 µL) were carried out in 100 mM potassium phosphate buffer (pH 7.4) containing 3.0 mM $MgCl_2$ and 1.0 mM β-NADPH at 37°C for 0 and 60 minutes. Following incubation, reactions were quenched with 800 µL of acetonitrile, centrifuged, and the resulting supernatants were evaporated under nitrogen. Dried residues were reconstituted in 200 µL of 30% acetonitrile and analyzed via LC-UV-MS.

LC-UV-MS Conditions:

Chromatographic analysis of **MP-004** was conducted using a Shimadzu HPLC system with an ACE 3 C18-AR column (150 × 4.6 mm) maintained at 30°C and equipped with a PDA detector (λ = 190–400 nm). The mobile phase consisted of (A) 0.1% formic acid and 2 mM ammonium formate in water/acetonitrile (95:5, v/v) and (B) 0.1% formic acid and 2 mM ammonium formate in water/acetonitrile (5:95, v/v). A 15 µL injection volume was used, and separation was achieved with a gradient elution at a flow rate of 700 µL/min over 55 minutes. Mass spectrometric detection was performed using a Thermo QExactive instrument in positive electrospray ionization mode, with spray voltage at 3.5 kV, capillary temperature at 350°C, and gas pressures set at 60 (sheath), 20 (auxiliary), and 10 (ion sweep) units. Data acquisition was performed in Q1 and MS2 scan modes using Xcalibur software.

Analysis of **7-ethoxycoumarin** was carried out using a Shimadzu UFLC-20A system coupled to a PDA detector (λ = 190–400 nm) and an ACE 3 C18-AR column (150 × 4.6 mm) held at 35°C. The mobile phases were identical to those used for MP-004, with a flow rate of 700 µL/min and an injection volume of 20 µL. A gradient program over 35 minutes facilitated the separation. Detection was performed with a Thermo LTQ Orbitrap mass spectrometer in positive electrospray ionization mode, using a spray voltage of 3.5 kV, a capillary temperature of 350°C, and sheath, auxiliary, and ion sweep gas pressures of 50, 15, and 15 units, respectively. Q1 scan mode was used for data acquisition using Xcalibur software.

**Cytochrome P450 (CYP) Inhibition in Human Liver Microsomes:**

CYP inhibition studies were performed in 96-well plates. Each reaction well received 20 µL of phosphate buffer added to blank wells. Then, 2 µL of test compound or positive control were added to their respective wells, and 2 µL of solvent was added to no-inhibitor and blank wells. Human liver microsomes were added to each well, followed by pre-incubation for 10 minutes at 37°C. After preparing the NADPH cofactor solution, 20 µL was added to each well to initiate the reaction, which continued for 10 minutes at 37°C. The reactions were stopped by adding 400 µL of cold stop solution (200 ng/mL tolbutamide and labetalol in acetonitrile). Samples were centrifuged at 4000 rpm for 20 minutes, and 200 µL of supernatant was mixed with 100 µL of HPLC-grade water and shaken for 10 minutes. Samples were analyzed by LC-MS/MS.

**Plasma Stability:**

Pooled human plasma was centrifuged at 4000 rpm for 5 minutes to remove clots, and aliquoted (98 µL/well) into 96-well plates using an Apricot automated workstation. The test compound (2 µL of 100 µM) was added to wells designated for time points (T0, T10, T30, T60, T120), while blank wells received no compound. Plates were incubated at 37°C, and at designated time points, reactions were quenched by adding 400 µL of cold stop solution (200 ng/mL tolbutamide and 200 ng/mL labetalol in acetonitrile) to precipitate proteins. Plates were sealed, shaken for 20 minutes, and centrifuged at 4000 rpm at 4°C for 20 minutes. Supernatants (50 µL) were transferred into 100 µL HPLC-grade water, sealed, shaken for 10 minutes, and analyzed by LC-MS/MS.

**Dissociation constant (pKa) Determination:**

To determine the aqueous pKa, ionic strength-adjusted water (ISA water, 0.15 M KCl) and 80% v/v methanol (MeOH) with 0.15 M KCl were prepared. For UV-metric pKa determination, 5 µL of 10 mM sample stock in DMSO and 25 µL of UV buffer were added to a vial, followed by 1.5 mL of 80% MeOH. For pH-metric pKa determination, ~1 mg of the test compound was weighed directly into the vial and 1.5 mL of 80% MeOH was added. In both assays, the instrument automatically pre-acidified the solutions with 0.5 M HCl and performed three titrations from low to high pH. Resulting curves were analyzed to determine the aqueous pKa value.

**Aqueous solubility:**

The aqueous solubility of test compound was determined using 0.1 N HCl and 50 mM phosphate buffer (PB) at pH 3.5, 4.5, and 7.4. Buffers were prepared by combining appropriate ratios of 50 mM $H_3PO_4$, $NaH_2PO_4$, and $Na_2HPO_4$ to achieve the target pH values. For each condition, 10 µL of a 10 mM DMSO stock solution of the test or control compound was added to the lower chamber of Whatman Mini-UniPrep vials, followed by 490 µL of buffer. Samples were vortexed for 2 minutes and shaken at RT for 24 hours at 800 rpm. After incubation, samples were centrifuged at 4000 rpm for 20 minutes. The vials were then compressed to filter the supernatants, which were analyzed by UPLC. Final concentrations were calculated using a standard calibration curve.

**Partition Coefficient Determination:**

Partitioning experiments were conducted using phosphate buffers (100 mM) at pH 1.8, 2.5, 4.5, 7.4, 9.5, 11, and 12 prepared by mixing $NaH_2PO_4$, $Na_2HPO_4$, $H_3PO_4$, and 5 N NaOH as needed. To obtain phase-saturated solvents, 10 mL of 1-octanol were mixed with 100 mL of each buffer to prepare octanol-saturated buffer, while in parallel, 10 mL of each buffer were mixed with 100 mL of 1-octanol to prepare buffer-saturated octanol.

Mixtures were shaken vigorously and equilibrated overnight at RT. For each condition, 2 µL of a 10 mM DMSO stock solution of the test compound was added to tubes in duplicate, followed by 149 µL of buffer-saturated octanol and 149 µL of octanol-saturated buffer. Tubes were vortexed for 2 minutes and shaken at 800 rpm for 1 hour at RT. Samples were then centrifuged at 4000 rpm for 5 minutes, and both aqueous and organic phases were collected and diluted as needed. Compound concentrations were quantified by LC-MS/MS using an ACQUITY UPLC BEH C18 column (1.7 µm, 2.1 × 50 mm) with 0.1% formic acid in water (mobile phase A) and 0.1% formic acid in acetonitrile (mobile phase B). Log D values were calculated from the ratio of compound concentrations in the octanol and buffer phases.

**Evaluation of effects on hERG current in stably transfected HEK-293 cells**

A concentration response relationship was established for the test item in 3 stably transfected HEK-293 cells. On each cell, the following treatments were tested: Tyrode's solution, vehicle of MP-004 (Tyrode's solution), MP-004 at $9x10^{-7}$ M, $9x10^{-6}$ M and $9x10^{-5}$ M. The method control item (E-4031) was tested at 3 concentrations (1, 10 and 100 nM) in one separate HEK-293 cell to support the validity of the method. The cells were exposed to the test item for at least 5 minutes. An $IC_{50}$ value was determined after observed more than 30% inhibition of hERG tail current was observed at the highest concentration.

Cells were clamped at -80 mV, depolarized to 0 mV for 5 sec allowing activation of hERG current and repolarized to -50 mV for 5 sec allowing hERG tail current to deactivate. This procedure was repeated at a frequency of 0.067 Hz. The current amplitude upon repolarization (hERG tail current) was measured before and after exposure to the test item. Experiments were conducted and currents acquired by means of pClamp software.

#### SUPPLEMENTARY FIGURES

**Supplementary Figure 1.** MP-004 off-targets safety panel.

| Target | IC50 (nM) | MaxDose (nM) | %Inh@Max Dose | Reference | MP-004 |
|---|---|---|---|---|---|
| | | | | | 10 µM |
| Apha2A | 8.50 | 1000 | 99.88 | yohimbine | 13.85 |
| 5HT3 | 38.53 | 10000 | 99.92 | MDL 72222 | 9.53 |
| NMDA | 10.79 | 10000 | 97.74 | MK-801 | -0.72 |
| CCKa | 0.96 | 1000 | 99.23 | CCK-8s | 3.09 |
| 5HT1B | 67.63 | 10000 | 100.71 | serotonin | 6.67 |
| CB1 | 1.35 | 1000 | 103.01 | CP 55940 | 1.31 |
| CB2 | 1.68 | 1000 | 103.76 | WIN 55212-2 | 28.11 |
| D1 | 0.73 | 1000 | 101.27 | SCH 23390 | 1.91 |
| D2 | 3.02 | 1000 | 102.36 | 7-OH-DPAT | 16.20 |
| hERG | 2.79 | 1000 | 93.89 | dofetilide | 34.79 |
| 5HT2B | 55.27 | 10000 | 101.09 | (±)DOI | 35.05 |
| M2 | 3.52 | 1000 | 100.92 | methoctramine | 15.54 |
| M3 | 1.49 | 1000 | 101.49 | 4-DAMP | 3.06 |
| 5HTT | 1.99 | 1000 | 111.85 | imipramine | -26.02 |
| NET | 3.52 | 1000 | 102.88 | protriptyline | -6.89 |
| ADORA2A | 1.30 | 1000 | 105.42 | CGS 15943 | -8.20 |
| GABAA | 1.70 | 1000 | 101.68 | flumazenil | -8.64 |
| H1 | 4.21 | 1000 | 99.09 | pyrilamine | 34.86 |
| M1 | 6.61 | 1000 | 97.78 | pirenzepine | 10.86 |
| 5HT2a | 2.18 | 1000 | 100.21 | Ketanserin | 24.68 |
| DAT | 16.82 | 1000 | 105.28 | BTCP | -19.01 |
| Ca2+-L | 0.23 | 1000 | 104.34 | nitrendipine | 2.46 |
| Apha1A | 0.60 | 100 | 99.72 | WB 4101 | 2.97 |
| 5HT1A | 0.73 | 1000 | 94.35 | 8-OH-DPAT | 2.33 |
| H2 | 566.80 | 10000 | 96.88 | cimetidine | 19.61 |
| op-delta | 0.22 | 100 | 106.07 | naltrindole | 1.70 |
| op-mu | 0.69 | 1000 | 100.96 | DAMGO | 8.57 |
| Beta1 | 652.90 | 100000 | 87.15 | atenolol | -4.50 |
| Beta2 | 1.28 | 1000 | 97.63 | ICI 118551 | -1.42 |
| nACHR-alpha7 | 27.46 | 1000 | 97.97 | MLA | 7.50 |
| op-kappa | 4.20 | 1000 | 99.31 | U-50488 | 1.38 |
| V1a | 0.36 | 100 | 102.64 | [d(CH2)51,Tyr(Me)2]-AVP | 9.95 |
| AR | 10.12 | 1000 | 100.26 | Progesterone | -3.92 |
| MAO-A | 5.37 | 1000 | 100.05 | Clorgyline | -4.31 |
| ACHE | 125.30 | 100000 | 96.45 | Neostigmine bromide | 2.56 |
| COX1 | 21.17 | 5000 | 99.17 | Diclofenac | -3.00 |
| COX2 | 640.50 | 50000 | 101.53 | NS-398 | -2.93 |
| PDE3A | 0.26 | 50 | 99.86 | Trequinsin hydrochloride | 15.87 |
| PDE4D2 | 17570 | 1000000 | 100.05 | IBMX | -10.60 |
| LCK | 0.49 | 200 | 139.24 | stausporine | 8.63 |

| Target | EC50 (nM) | Max dose nM | Act%@ max dose | Reference ID | MP-004 |
|---|---|---|---|---|---|
| | | | | | 12 µM |
| Eta Agonist | 1.14 | 300 | 103.00 | Endothelin 1 | -0.10 |

| Target | IC50 (nM) | Max dose nM | Inh%@ max dose | Reference ID | MP-004 |
|---|---|---|---|---|---|
| | | | | | 10 µM |
| Eta Antagonist | 2.67 | 75 | 100.32 | BQ-123 | 0.78 |

| hNav1.5 | | | | | |
|---|---|---|---|---|---|
| Compound | Conc (µM) | % Inhibition | | Average | Std Deviation |
| Tetracaine | 10 | 99.36 | - | 99.36 | - |
| MP-004 | 10 | 7.70 | 3.39 | 5.55 | 2.15 |

| hKCNQ1 | | | | | |
|---|---|---|---|---|---|
| Compound | Conc (µM) | % Inhibition | | Average | Std Deviation |
| Chromanol 293B | 10 | 41.19 | - | 41.19 | - |
| MP-004 | 10 | 15.38 | 22.55 | 18.96 | 3.58 |

**Supplementary Figure 2.** Bidirectional permeability in Caco-2 cells.

| Compound ID | Mean $P_{app}$ ($10^{-6}$ cm/s) | | Efflux Ratio | Mean %Solution Recovery | | Rank | | Note |
|---|---|---|---|---|---|---|---|---|
| | A to B | B to A | | A to B | B to A | $P_{app}$ | Efflux Transporter Substrate | |
| Nadolol | 0.097 | ND | - | 96 | ND | Low | - | Low permeability marker |
| Metoprolol | 20.1 | ND | - | 102 | ND | High | - | High permeability marker |
| Digoxin | 0.0334 | 14,6 | 437 | 96.5 | 98.4 | Low | Likely | P-gp substrate |
| MP004 | 40.2 | 31.8 | 0.79 | 90.1 | 94 | High | Poor or non | - |

ND: not determined

Binning Criteria:

- Low permeability: $P_{app} \leq 0.500$ ($\times 10^{-6}$ cm/s)
- Moderate permeability: $0.500 < P_{app} < 2.50$ ($\times 10^{-6}$ cm/s)
- High permeability: $P_{app} \geq 2.50$ ($\times 10^{-6}$ cm/s)

Substrate Potential:

- Likely: ER ≥ 2.00
- Poor or non: ER < 2.00

**Supplementary Figure 3.** P-gp substrate assessment for MP-004 in the Caco-2 cells.

| Compound ID | Inhibitor | Mean $P_{app}$ ($10^{-6}$ cm/s) | | Efflux Ratio | Mean % Solution Recovery | | Rank | | | Note |
|---|---|---|---|---|---|---|---|---|---|---|
| | | A to B | B to A | | A to B | B to A | $P_{app}$ | Efflux Transporter Substrate | P-gp Substrate | |
| Nadolol | / | 0.09 | ND | - | 96.5 | ND | Low | - | - | Low permeability marker |
| Metoprolol | / | 20.1 | ND | - | 101.6 | ND | High | - | - | High permeability marker |
| Digoxin | / | 0.03 | 14.6 | 437 | 96.5 | 98.4 | Low | Probably | Probably | P-gp substrate |
| | Verapamil | 1.31 | 9.78 | 7.44 | 87.6 | 97.3 | | | | |
| MP-004 | / | 40.2 | 31.8 | 0.79 | 90.1 | 94.0 | High | Poor or non | Poor or non | - |
| | Verapamil | 43.2 | 34.2 | 0.79 | 111.2 | 123.7 | | | | |

ND: not determined

Binning Criteria:

- Low permeability: $P_{app} \leq 0.5$ ($\times 10^{-6}$ cm/s)
- Moderate permeability:$0.5 < P_{app} < 2.5$ ($\times 10^{-6}$ cm/s)
- High permeability: $P_{app} \geq 2.5$ ($\times 10^{-6}$ cm/s)

Substrate Potential:

- Probably: $ERa \geq 2$ and $ERa / ER_i > 2$
- Likely: $ERa \geq 2$
- Poor or non: $ERa < 2$

**Supplementary Figure 4.** MP-004 plasma proteins binding.

| Compound ID | Species / Matrix | %Unbound | % Unbound SD | %Bound | %Recovery |
|---|---|---|---|---|---|
| MP-004 | CD-1 Mouse Plasma | 47.52 | 3.8 | 52.48 | 100.7 |
| | SD Rat Plasma | 33.31 | 4.3 | 66.69 | 103.3 |
| | NZW Rabbit Plasma | 47.9 | 9.9 | 52.1 | 105.4 |
| | Beagle Dog Plasma | 47.61 | 0.3 | 52.39 | 109 |
| | Göttingen minipig Plasma | 50.2 | 2.2 | 49.8 | 116.9 |
| | Cynomolgus Monkey Plasma | 42.28 | 6.5 | 57.72 | 115.8 |
| | Human Plasma | 37.75 | 2.4 | 62.25 | 91.8 |
| Warfarin | CD-1 Mouse Plasma | 4.17 | 0.5 | 95.83 | 103.5 |
| | SD Rat Plasma | 0.77 | 0.1 | 99.23 | 105.4 |
| | NZW Rabbit Plasma | 4.8 | 0.9 | 95.2 | 95.3 |
| | Beagle Dog Plasma | 4.18 | 0.6 | 95.82 | 102.6 |
| | Göttingen minipig Plasma | 1.6 | 0.1 | 98.4 | 97.3 |
| | Cynomolgus Monkey Plasma | 1.58 | 0.2 | 98.42 | 99.1 |
| | Human Plasma | 1.11 | 0.1 | 98.89 | 102.6 |

**Supplementary Figure 5.** Metabolite profiling in liver microsomes

| Metabolite Code | $[M + H]^+$ *m/z* | RT (min) | Relative Abundance (UV peak area %Total) | | | | | | Metabolic Pathways |
|---|---|---|---|---|---|---|---|---|---|
| | | | Mouse | Rat | Dog | Rabbit | Minipig | Human | |
| M1 | 295.12 | 12.92 | 2.04 | 7.91 | + | 4.83 | 9.95 | 2.59 | Mono-oxidation and demethylation (P+O –$CH_2$) |
| M2 | 309.14 | 13.18 | 14.28 | 17.29 | 2.74 | 16.68 | 3.98 | 19.90 | Mono-oxidation (P + O) |
| M3 | 325.13 | 13.55 | 1.60 | 6.44 | 2.35 | 5.29 | + | 1.87 | Di-oxidation (P + 2O) |
| M4 | 265.11 | 14.46 | 0.61 | 1.39 | 0.10 | 0.44 | + | 0.32 | Di-demethylation (P –2$CH_2$) |
| M5 | 279.13 | 14.97 | + | 2.76 | ND | 1.75 | + | 10.25 | Demethylation (P –$CH_2$) |
| M6 | 325.13 | 15.77 | 0.88 | 0.20 | 0.18 | 1.35 | + | + | Di-oxidation (P + 2O) |
| M7 | 295.12 | 15.95 | 0.58 | 4.08 | 0.45 | 0.90 | + | 2.99 | Mono-oxidation and demethylation (P+O –$CH_2$) |
| M8 | 265.11 | 21.59 | 0.75 | 1.51 | 0.77 | + | + | + | Di-demethylation (P –2$CH_2$) |
| M9 | 279.13 | 22.35 | 4.34 | 4.20 | 2.29 | 1.96 | + | 8.30 | Demethylation (P –$CH_2$) |
| M10 | 309.14 | 24.29 | 48.87 | 39.11 | 60.06 | 37.52 | 58.45 | 2.50 | Mono-oxidation (P + O) |
| MP-004 | 293.14 | 23.04 | 26.05 | 15.10 | 31.05 | 29.26 | 27.62 | 51.27 | NA |

Note: +: Only detected by LC-MS; P: parent drug.

$$\%\text{Total} = \frac{\text{Peak Area of a Related Component}}{\text{Peak Area of Total Related Components}} \times 100\%$$

**Supplementary Figure 6.** CYP inhibition in liver microsomes.

Results summary:

| Compound ID | $IC_{50}$ (μM) | | | | |
|---|---|---|---|---|---|
| | CYP1A2 | CYP2C9 | CYP2C19 | CYP2D6 | CYP3A4-M |
| MP-004 | >50 | >50 | 35.8 | 45.5 | 43.1 |

Positive controls:

| CYP Isozyme | Standard Inhibitor | %Inhibition | %Inhibition Acceptance Range | Pass/No Pass |
|---|---|---|---|---|
| CYP1A2 | α-Naphthoflavone | 88.8 | 82.1%-94.9% | Pass |
| CYP2C9 | Sulfaphenazole | 84.1 | 79.0%-89.9% | Pass |
| CYP2C19 | (+)-N-3-benzylnirvanol | 82.6 | 77.5%-91.3% | Pass |
| CYP2D6 | Quinidine | 96.1 | 93.9%-97.0% | Pass |
| CYP3A4 | Ketoconazole | 98.7 | 97.6%-99.9% | Pass |

**Supplementary Figure 7.** MP-004 stability in plasma.

| Compound ID | Time Point (min) | % Remaining | $T_{1/2}$ (min) |
|---|---|---|---|
| MP-004 | 0 | 100 | >289.1 |
| | 10 | 97.9 | |
| | 30 | 98.7 | |
| | 60 | 109.9 | |
| | 120 | 91.5 | |
| Propantheline bromide | 0 | 100 | 9.6 |
| | 10 | 60.2 | |
| | 30 | 13.6 | |
| | 60 | 1.4 | |
| | 120 | 0 | |

**Supplementary Figure 8.** MP-004 dissociation constant (pKa)

| Compound ID | Items | pKa (UV metric) | pKa (pH metric) | Final pKa Result | Comments |
|---|---|---|---|---|---|
| MP-004 | **pKa 1** | 7.09 | 7.27 | 7.18 | Final pKa was obtained by UV metric at pH 2-12 and pH metric method at pH 3-11 |
| | **pKa 2** | 11.15 | NA | 11.5 | |

**Supplementary Figure 9.** MP-004 aqueous solubility

| Compound ID | Media | Visual Appearance | Kinetic Solubility (µM) | Kinetic Solubility (µg/mL) |
|---|---|---|---|---|
| MP-004 | 0.1 N HCl | No visible particles | 228 | 75 |
| | PB (pH 3.5) | No visible particles | 229 | 75.4 |
| | PB (pH 4.5) | No visible particles | 228 | 75 |
| | PB (pH 7.4) | No visible particles | 216 | 70.9 |

**Supplementary Figure 10.** MP-004 partition coefficient MP-004 permeability.

| Compound ID | Log P (Oct/buff) | log D (pH 1.8) | log D (pH 2.5) | log D (pH 4.5) | log D (pH 7.4) | log D (pH 9.5) | log D (pH 11.0) | log D (pH 12.0) |
|---|---|---|---|---|---|---|---|---|
| MP-004 | 1.77 | -2.07 | -1.79 | -0.679 | 1.49 | 1.77 | 1.75 | 1.8 |

**Supplementary Figure 11.** Effect of MP-004 on hERG tail current amplitude.

| | | | MP-004 | | |
|---|---|---|---|---|---|
| **Cell** | **Tyrode** | **Vehicle** | **9x10$^{-7}$M** | **9x10$^{-6}$M** | **9x10$^{-5}$M** |
| Cell 1 | 924 | 4 | 6 | 8 | 44 |
| Cell 2 | 742 | 4 | 3 | 11 | 59 |
| Cell 3 | 670 | 3 | 3 | 12 | 57 |
| Mean | 779 | 4 | 4 | 10 | 53 |
| SEM | 76 | 0 | 1 | 1 | 5 |

**Supplementary Figure 12.** Serum, retina, optic nerve, brain, kidney and liver pharmacokinetic parameters of MP-004 in rabbits following ocular instillation.

| PK Parameters | Serum | Retina | Optic nerve | Brain | Kidney | Liver |
|---|---|---|---|---|---|---|
| $C_{max}$ (ng/mL or ng/g) | 43.6 | 387 | 158 | 42.2 | 145 | 16.7 |
| $T_{max}$ (h) | 0.5 | 1 | 1 | 1 | 1 | 1 |
| $T_{1/2}$ (h) | 0.3 | 6.3 | 4.81 | ND | ND | ND |
| $T_{last}$ (h) | 2 | 12 | 12 | ND | ND | ND |
| $AUC_{0\text{-}last}$ (ng.h/mL or ng.h/g) | 27.7 | 1460 | 441 | ND | ND | ND |
| $AUC_{0\text{-}24}$ (ng.h/mL or ng.h/g) | 28.2 | 1719 | 549 | ND | ND | ND |
| $AUC_{0\text{-}inf}$ (ng.h/mL or ng.h/g) | 28.2 | 1813 | 582 | ND | ND | ND |

"ND" means not determined.

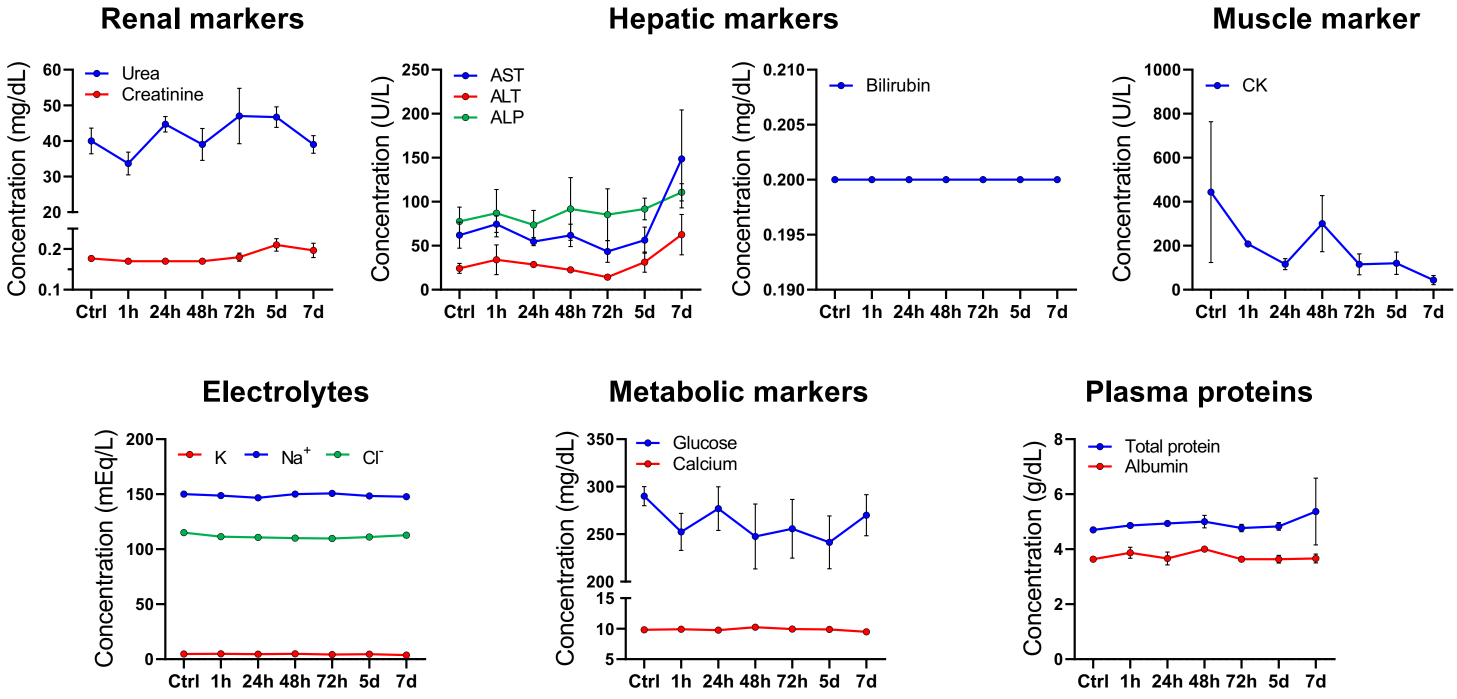


**Supplementary Figure 13. Analysis of the different biochemical parameters measured in mice serum.** Blood test was conducted in control (Ctrl) group and 1, 24, 48, 72 hours (h) and 5 and 7 days (d) after last-administration groups. Data are expressed as mean ± SEM. n=3 mice/time. One-way ANOVA statistical test followed by Dunnett multiple comparisons post-hoc test vs control (Ctrl, non treated) was performed. AST, aspartate aminotransferase; ALT, alanine aminotransferase; ALP, alkaline phosphatase; CK, creatinine kinase; K, potassium; $Na^+$, sodium; $Cl^-$, chloride.

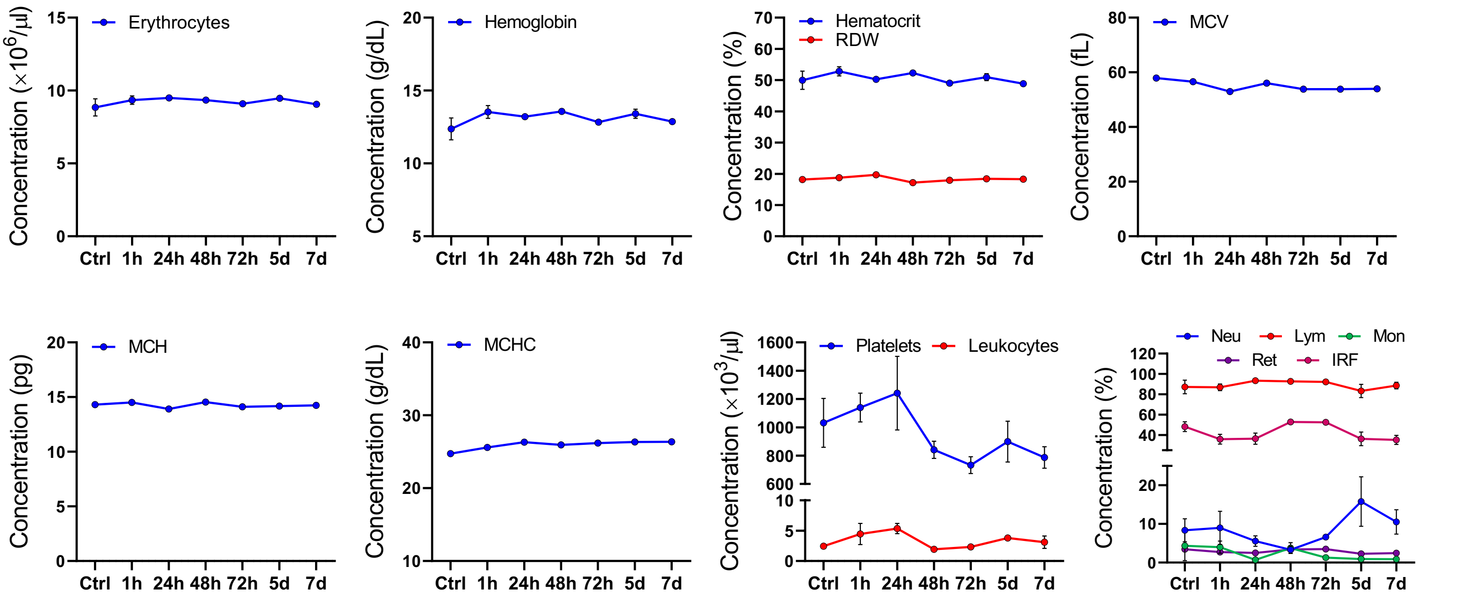


**Supplementary Figure 14. Analysis of the different hematological parameters measured in mice serum**. Blood test was conducted in control (Ctrl) group and 1, 24, 48, 72 hours (h) and 5 and 7 days (d) after last-administration groups. Data are expressed as mean ± SEM. n=3 mice/time. One-way ANOVA statistical test followed by Dunnett multiple comparisons post-hoc test vs control (Ctrl, non-treated) was performed. RDW, Red cell Distribution Width; MCV, mean corpuscular volume; MCH, erythrocyte mean corpuscular hemoglobin; MCHC, mean corpuscular hemoglobin concentration; Neu, neutrophils; Lym, lymphocytes; Mon, monocytes; Ret, reticulocytes; IRF, Immature Reticulocytes Fraction.

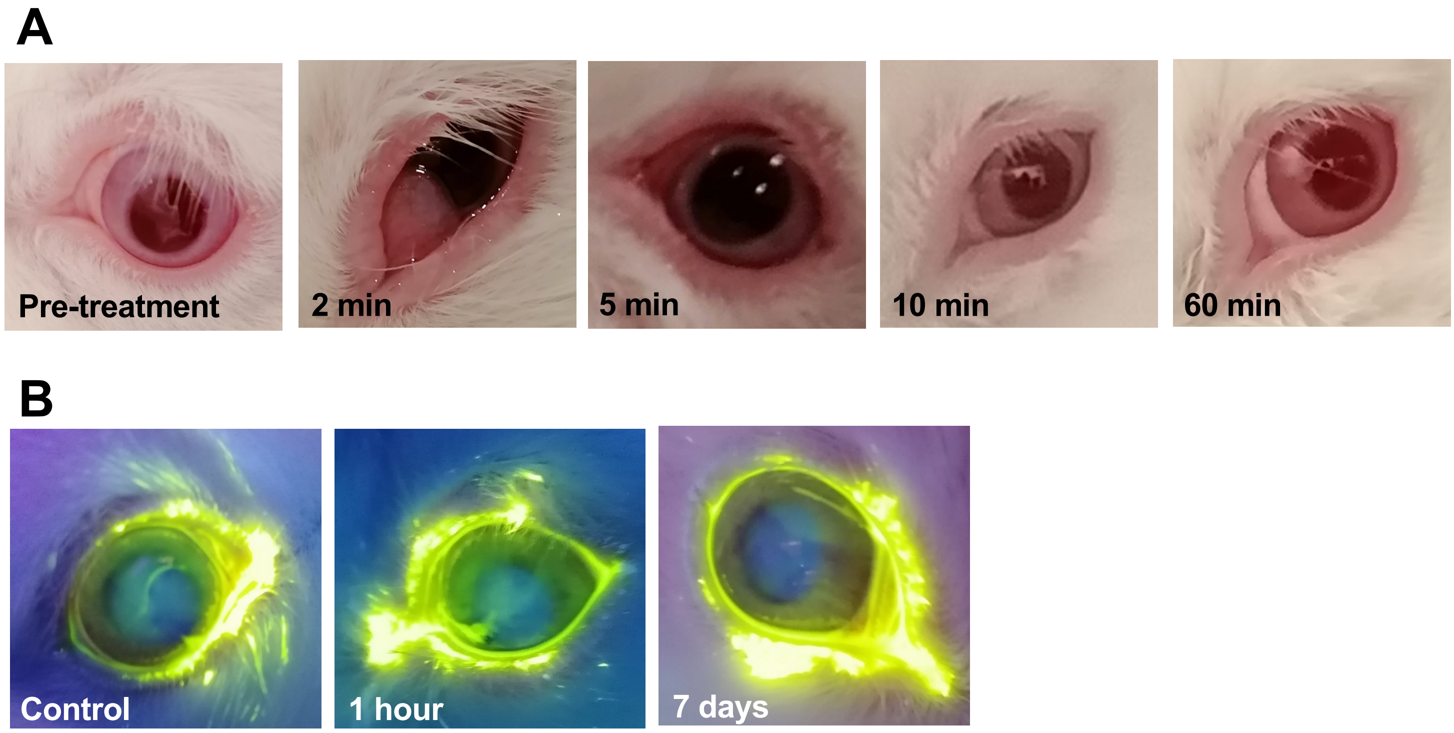


**Figure Supplementary 15. Findings in the ocular observations of the toxicological rabbit's assay. A)** Representative images of the transient redness produced after treatment with MP-004. Pre-treatment, 2, 5, 10 and 60 minutes after topical ocular administration are shown. **B)** Representative fluorescein images of rabbit eyes. Control (vehicle-treated) group, 1 hour and 7 days post-administration groups images are shown.

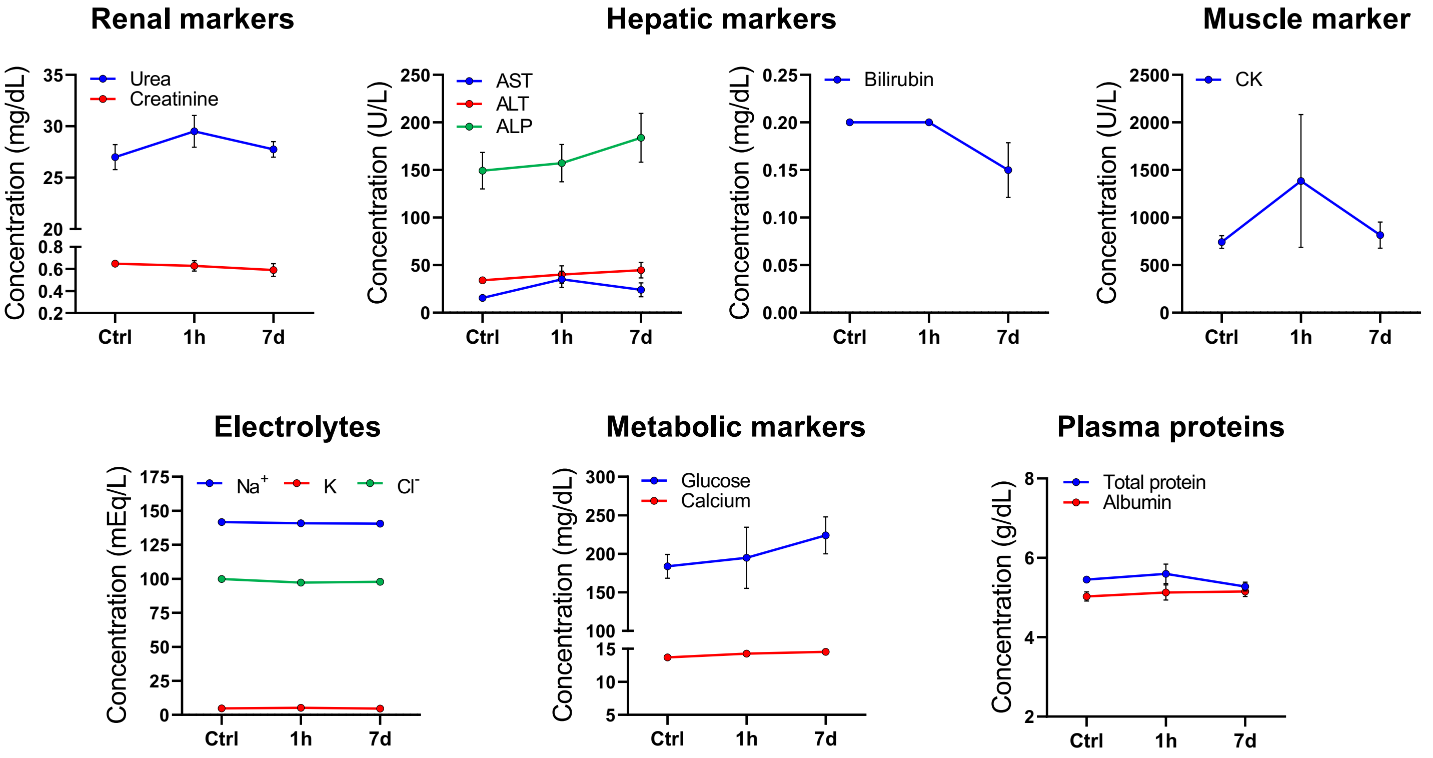


**Supplementary Figure 16. Analysis of the different biochemical parameters measured in rabbit serum.** Blood test was conducted in control (Ctrl) group and 1, 24, 48, 72 hours (h) and 5 and 7 days (d) after last-administration groups. Data are expressed as mean ± SEM. n=4 rabbits/time. One-way ANOVA statistical test followed by Dunnett multiple comparisons post-hoc test vs control (Ctrl, vehicle-treated) was performed. AST, aspartate aminotransferase; ALT, alanine aminotransferase; ALP, alkaline phosphatase. CK, creatine kinase

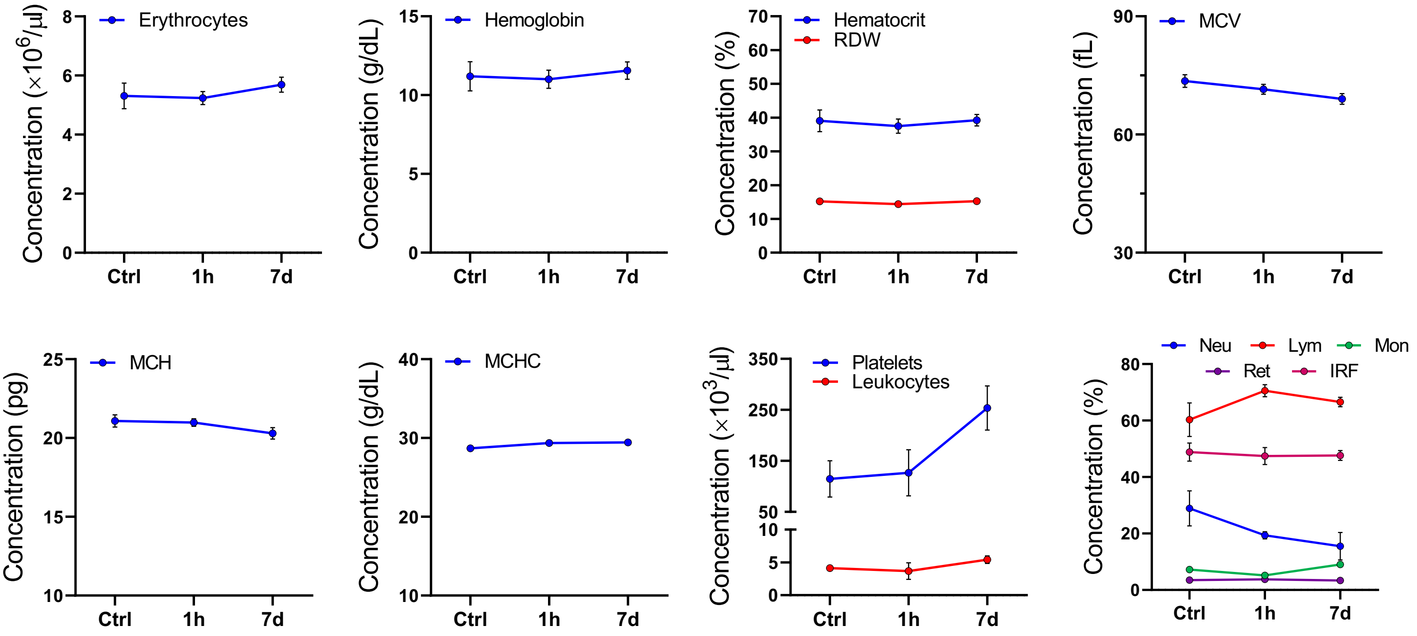


**Supplementary Figure 17. Analysis of the different hematological parameters measured in rabbit serum**. Blood test was conducted in control (Ctrl) group and 1, 24, 48, 72 hours (h) and 5 and 7 days (d) after last-administration groups. Data are expressed as mean ± SEM. n=4 rabbits/time. One-way ANOVA statistical test followed by Dunnett multiple comparisons post-hoc test vs control (Ctrl, vehicle-treated) was performed. RDW, Red cell Distribution Width; MCV, mean corpuscular volume; MCH, erythrocyte mean corpuscular hemoglobin; MCHC, mean corpuscular hemoglobin concentration; Neu, neutrophils; Lym, lymphocytes; Mon, monocytes; Ret, reticulocytes; IRF, Immature Reticulocytes Fraction.
